# Collective motion conceals fitness differences in crowded cellular populations

**DOI:** 10.1101/267286

**Authors:** Jona Kayser, Carl Schreck, Matti Gralka, Diana Fusco, Oskar Hallatschek

## Abstract

Many cellular populations are tightly-packed, for example microbial colonies and biofilms [39, 10, 41], or tissues and tumors in multi-cellular organisms [11, 29]. Movement of one cell inside such crowded assemblages requires movement of others, so that cell displacements are correlated over many cell diameters [28, 6, 31]. Whenever movement is important for survival or growth [15, 34, 38, 9], such correlated rearrangements could couple the evolutionary fate of different lineages. Yet, little is known about the interplay between mechanical stresses and evolution in dense cellular populations. Here, by tracking deleterious mutations at the expanding edge of yeast colonies, we show that crowding-induced collective motion prevents costly mutations from being weeded out rapidly. Joint pushing by neighboring cells generates correlated movements that suppress the differential displacements required for selection to act. Such mechanical screening of fitness differences allows the mutants to leave more descendants than expected under non-mechanical models, thereby increasing their chance for evolutionary rescue [2, 5]. Our work suggests that mechanical interactions generally influence evolutionary outcomes in crowded cellular populations, which has to be considered when modeling drug resistance or cancer evolution [1, 22, 34, 30, 36, 42].

---

As a model system for crowded cellular populations, we focused on colonies of the budding yeast *Saccharomyces cerevisiae* [33]. Since yeast cells lack motility, colony expansion is fueled purely by the pushing forces generated by cellular growth and division [14, 7]. To explore how these pushing forces affect the strength of natural selection, we competed “mutant” cells, carrying a growth-rate deficit, with faster-growing “wild-type” cells in expanding colonies (Fig. 1a–c). Mutant and wild-type cells remained in well-segregated sectors during the expansion [15], which allowed us to use time-resolved fluorescence microscopy to monitor the gradual demise of the mutant fraction. We then measured the rate at which mutant cells are out-competed by faster-growing wild-type cells at the expanding front of a linear colony (Fig. 1c).

We found that mutants were weeded out from the expanding frontier in two main stages. In the first stage, the width of mutant sectors decreased at a constant rate. This is in line with a minimal model where local front expansion velocities depend only on the cell-type specific growth rates of the “pioneer” cells at the leading edge [16] (dashed black lines in Fig. 1c, see SI Sec. 3 for model details). This “Local-speed” model predicts that the rate of decline of the mutant width remains constant until all mutants are expelled from the frontier, in agreement with different types of commonly-used non-mechanical simulations (SI Fig. 17 b, c, d). In contrast, we observed in our experiments a progressive slowdown of the rate of mutant decline as the width of mutant clones fell below about 200 μm. As a consequence, expulsion of the mutant clone from the expanding front was substantially delayed (yellow arrow in Fig. 1c). Evaluating the lengths of 41 independent clones revealed them to consistently exceed the null expectation by a substantial margin (Fig.1d).

**Figure 1:**
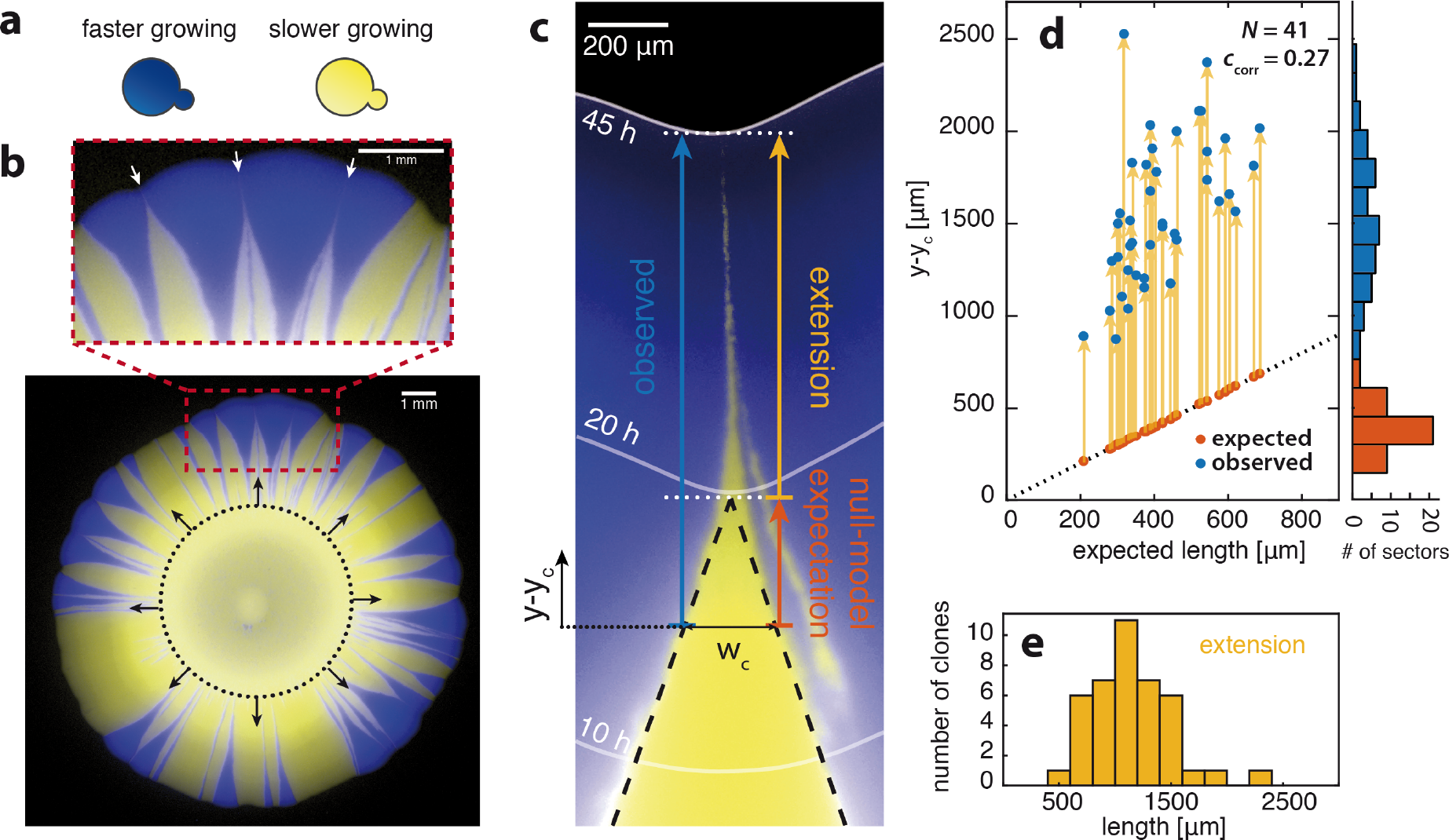
Attenuation of fitness differences in crowded microbial communities. a) Experiments are carried out by competing two types of yeast cells. In the presence of the drug cycloheximide yellow-colored cells have a growth rate disadvantage with respect to blue-colored cells (see SI Sec. 1 for details) [14]. b) Colonies grown from a mixture of both cell types (initial ratio of blue to yellow *r* = 10%, 80 nM cycloheximide). The black dotted circle indicates the area of initial cell deposition. As cells in the colony interior (bottom) do not grow, positions of mutant and wild-type cells are “frozen” in time as the front passes (solid gray lines mark the front position at the indicated times after inoculation) and represent a “fossil record” of past wild-type/mutant boundary positions. Zoom-in: Sectors of yellow cells caught between two expanding blue sectors exhibit elongated, funnel-shaped tips (white arrows). c) Fluorescent micrograph of a slower growing yellow sector (“mutant”) competing with adjacent blue sectors (“wild-type”). To avoid the additional geometric complexity of circular colonies, we initiated colony growth from a linear front in which sections of mutant cells were flanked by regions of wild-type cell (see SI Fig. 5). Initially, the clone boundaries converge at a constant angle set by their relative fitness (*s* = 0.06, 90 nM cycloheximide)[19, 14], in agreement with the Local-speed model (dashed black lines, see main text). As the width of the clone shrinks below about *w*_c_ = 230 μm, the boundaries begin to diverge from their initial linear trajectory, thereby delaying boundary collision. The observed length of the clone (blue arrow) after shrinking below the critical width *w*_c_ is substantially extended (yellow arrow) in comparison to the expected length (orange arrow) predicted by the Local-speed model (see supplementary movie S1). d) Expected and observed clone lengths for *N* = 41 individual sectors as a function of their expected length. All sectors show a significant (*p*-value « 10^−6^) extension of their lengths (yellow arrows) with the extension only weakly correlating with the expected sector lengths (*c*_corr_ = 0.27). The distributions of expected and observed lengths are shown as histograms (right). e) Histogram of the sector length extensions (yellow arrows in d)).

The fate of clones in crowded populations fundamentally depends on the relative motion of mutant and wild-type [14, 16, 35, 23, 19, 20]. The Local-speed model predicts that the direction of motion of pioneer cells at the mutant-wild type interface changes discontinuously. A delay of the extinction of mutant clones could arise from a suppression of such discontinuities. Indeed, while pioneer cells much further apart than 200 μm can have substantially different velocity vectors, their direction of motion becomes highly correlated as they approach one another (Fig. 2a). This directional alignment prevents the rapid collision of clone boundaries as they come close (dashed lines in Fig. 2a).

Since the average motion of pioneer cells is perpendicular to the local front (see Fig. 3b and SI Fig. 7), the directional alignment must be driven by a mechanism that suppresses high curvatures of the front line. Indeed, while the Local-speed model predicts that the front line develops a sharp kink (infinite curvature) as the mutant clone goes extinct (Fig 2a, orange frame), we find that the curvature of the front line saturates as the clone becomes smaller (Fig. 2a, compare 47 h and 62 h). Curvature suppression is an active, growth-driven process: Fig. 2b shows how an initial kink in the front, deliberately produced by initial cell deposition, is smoothed out by subsequent growth.

We hypothesized that the dynamic suppression of high front curvatures, and thus the directional alignment of nearby cells, could be described by an effective surface tension [26, 24, 3, 37]. To test this, we set up a phenomenological “Interface-growth” model where the population front moves along its local normal with a velocity that depends not only on cell-type but also on the local curvature of the front mediated by an effective surface tension (Fig. 2c and SI Sec. 5). The surface tension term dampens the growth of bulges and gives a boost to depressed regions of the front, as observed in our experiments (Fig. 2b). This model, which corresponds to a multitype extension of the so-called KPZ interface growth model [18] (see SI Sec. 5.3), reproduces the observed clone size dynamics very well upon tuning the line tension, our single fitting parameter (see Fig. 2d).

So, the attenuation of natural selection in our experiments can be traced back to the collective motion of nearby cells, which is driven by an effective surface tension that suppresses high front curvatures. This leads to the directional alignment of the motion of mutant pioneer cells with that of the flanking wild-type population, thereby concealing the fitness difference between both types. We can describe this screening effect by an effective selection difference between mutants and wild type that is sharply diminished for small clones, see Fig. 2e.

Which mechanism could generate an effective surface tension capable of driving the observed directional alignment? Since we expected mechanical cellcell forces to be the root cause for the collective cellular motion (see Fig. 3a, b), we tested *in silico* whether the required surface tension in a crowded population could be generated purely through the steric interactions between cells. To this end, we simulated growing and dividing cells as proliferating elastic spheres without any attractive interactions (Fig. 3e). To account for spatially heterogeneous growth rates, cells were allowed to grow only within a certain growth layer, consisting of all cells within a fixed distance from the edge of the colony (SI Sec. 6). We found that curved fronts gradually flattened as observed in colony experiments and predicted by our Interface-growth model (SI Fig. 5). In our cell-based simulations, the suppression of high curvatures can be rationalized by considering the behavior of a growth layer bent at constant curvature (Fig. 3d,e): If the growth layer is indented, more cells per unit length of front contribute to growth and pushing, thus leading to a local speed-up of the colony expansion. Conversely, we observe a local slow-down if the front exhibits a bulge so that less cells per unit length of front contribute to pushing. Thus, a negative feedback on front curvature, which is the basis for the observed velocity correlations across distances of many cell diameters, spontaneously arises from purely passive, mechanical effects. Finally, in our mechanical simulations, domain boundaries move perpendicular to the local front line (Fig. 12), as we had observed in our experiments and assumed in our Interface-growth model (see Fig. 2). This condition arises in any overdamped system where cell motion is driven by pressure gradients, since the edge of the colony represents a constant pressure line. It is violated, however, in common non-mechanical simulations of colony growth where domain boundaries are generically tilted with respect to the front line (SI Fig. 13). Our cell-based simulations thus generate the key ingredients of the Interface-growth model, an interface with a natural line tension and domain boundaries that move normal to the interface. Consequently, upon directly simulating our competition experiments, we reproduce funnel-like mutant clones that, near extinction, are carried along by the flanking wild-type cells (Fig. 3f).

**Figure 2:**
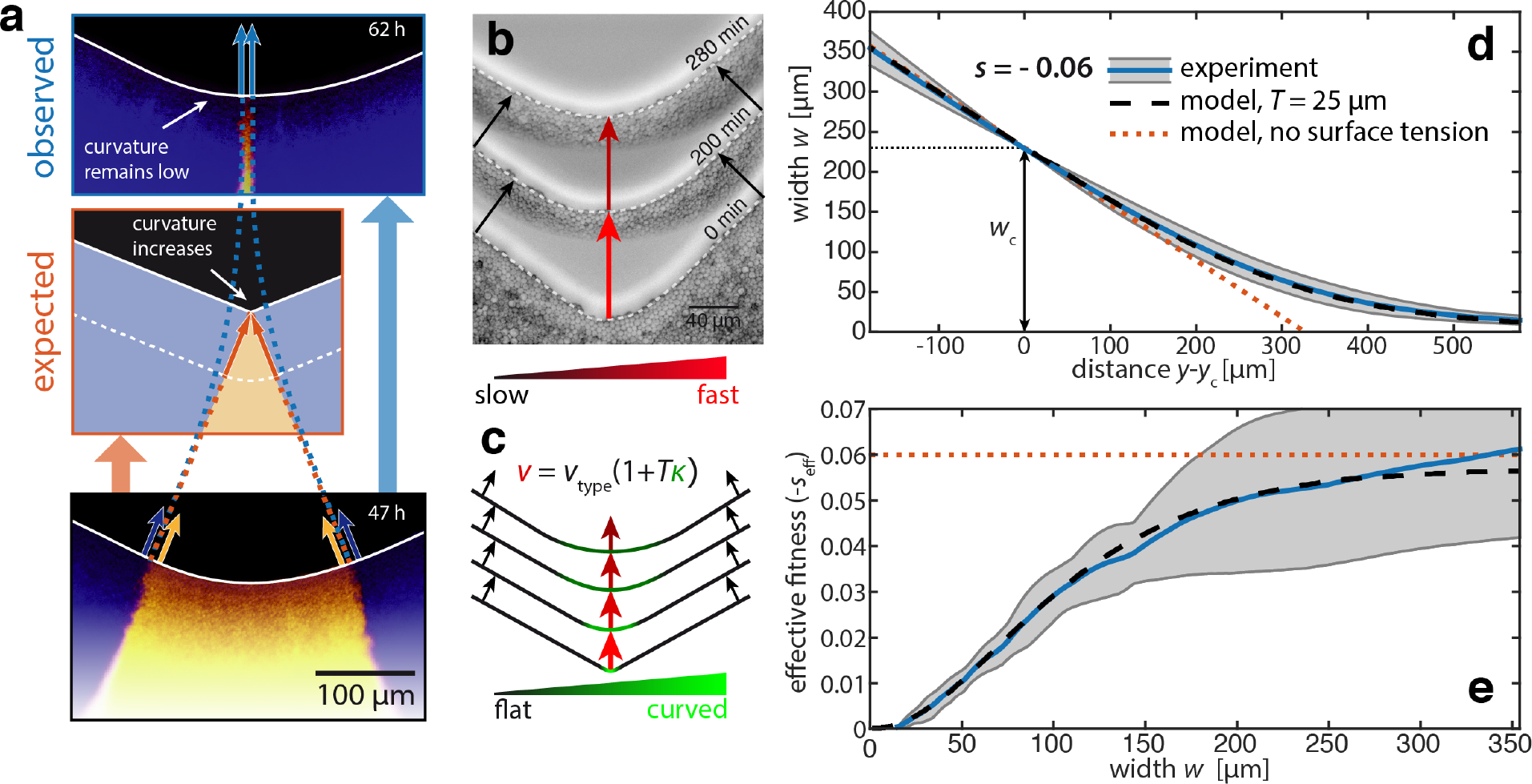
Directional alignment of clone-boundaries is due to an effective surface tension. a) Bottom (47h): Wild-type/mutant boundaries propagate perpendicular to the leading edge (solid white line, see SI Fig. 7) initially following straight, approaching trajectories (compare Fig. 1c) with wild-type and mutant cells at both sides of each boundary moving in a correlated manner (blue and yellow arrows). The front at the boundaries is almost flat while the mutant front between the boundaries is slightly curved. Center (orange, schematic): If boundaries were to remain on their straight path (dashed orange lines) the mutant front would have to continually increase in curvature, eventually forming a sharp kink just prior to boundary collision (orange arrows). Top (blue, 62h): Instead, we find that the low-curvature shape of the font is maintained. As a consequence, the trajectories of the two boundaries, following to the local front normal, have to align as they approach (dashed light-blue lines and light-blue arrows). b) Consecutive snap shots of a progressing front starting from cells deposited in a highly curved geometry. As expansion progresses, the front is rapidly flattened out due to increased displacement of the curved sections with respect to adjacent flat regions (red and black arrows, respectively). c) Schematic of Interface-growth model. Arrows indicate displacement of front between time steps (color code as in b)). Increased local curvature *κ* (green) causes a speed-up of the front proportional to the surface tension parameter *T*. *u*_type_ is equivalent to the front velocity in the Local-speed model. d) Clone width *w* (distance between boundaries) decays as a function of front position *y* (see Fig. 1c for definition of *y*_c_). The Interface-growth model (black dashed line) is fitted to the experimental data (solid blue line, gray shaded area indicates one standard deviation from experimental average, *N* = 9 independent sectors), yielding a surface tension of *T* = 25 μm. e) For each width *w*(*y*) one can define an effective fitness difference *s*_eff_(*w*) which decreases with declining clone width (see SI Sec. 3 for details). Labels as in d).

**Figure 3:**
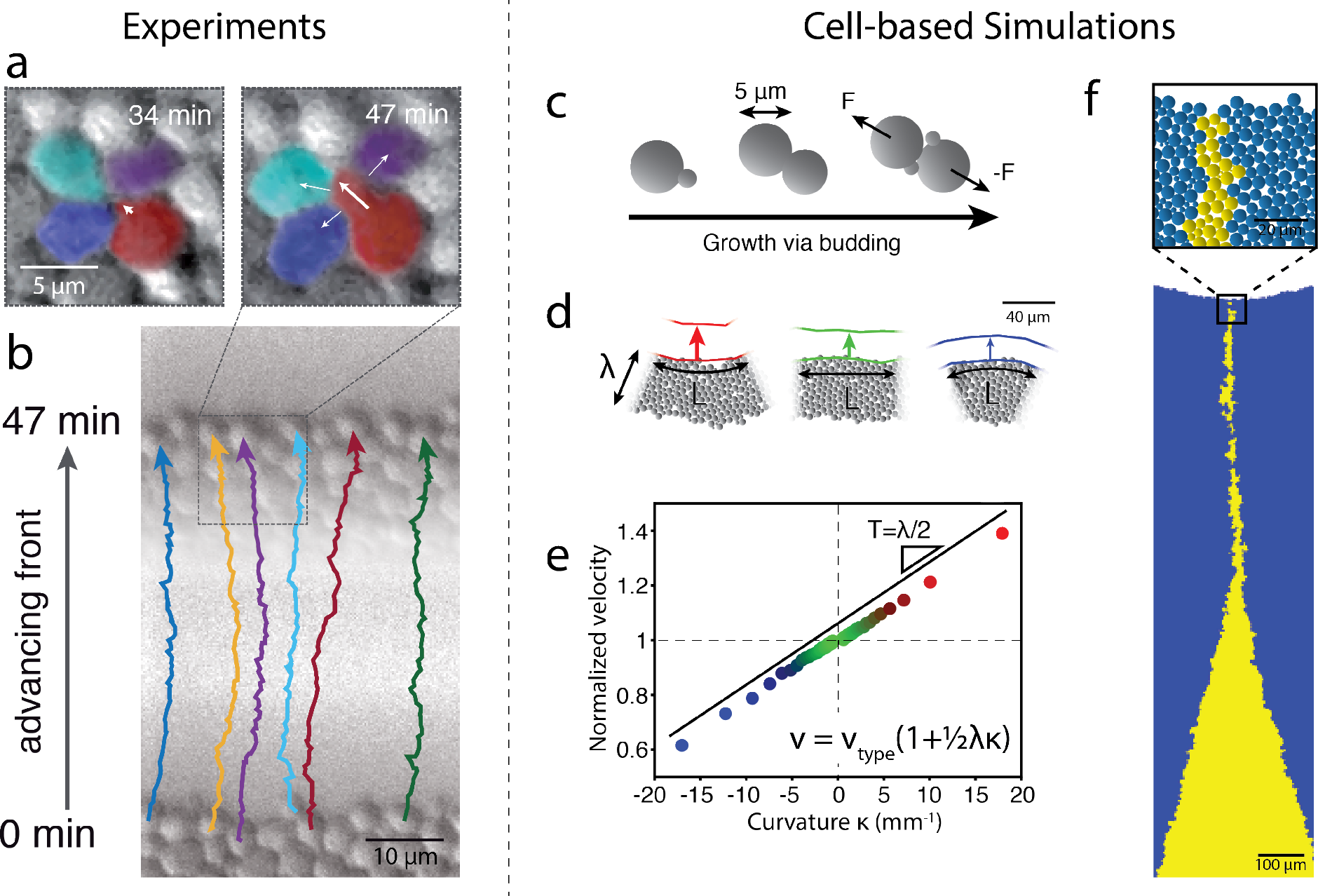
Effective surface tension emerges from purely steric cell-cell interactions. a) Yeast cells within a densely packed colony have to push their neighbors in order to proliferate. The thick arrow indicates the growth of a single budding cell (red) and thin arrows illustrate resulting forces acting on adjacent cells. b) Colored lines indicate tracks of individual cells at the expanding front that do not bud between the depicted time points (47 min, 1/2 generation). Nonetheless, these cells are displaced about 10 cell diameters, highlighting that cells move primarily due to cumulated pushing forces exerted by cells growing behind them, rather than their own growth. Due to symmetry, force components parallel to the front largely cancel each other out resulting in cells moving almost exclusively perpendicular to the front. c) Cell-based simulations model cells as conjoined spheres that proliferate via budding. Cells interact with repulsive contact forces and move according to overdamped dynamics. d) Cell growth occurs only in a “growth layer” of depth λ = 45 pm near the front. The front velocity is modulated by local front curvature (compare colored arrows). Indentations (left, red) move more quickly than flat fronts (center, green) due to a greater number of pushing cells per unit arc-length *L*. For the same reason bulges (right, blue) experience a slow down. Front profiles are shown for time-points separated by 1/2 generation. e) The velocity at the front increases linearly with front curvature k with linear coefficient (surface tension) *T*: *v* ∝ 1 + *Tκ*. Colors (blue to red) correspond to curvature and are the same as in d. f) Clones in cell-based simulations exhibit a funnel-like sector shape and delay in extinction, similar to those observed in experiments. Just prior to extinction, the mutant clone is extremely thin (≈ 10*μm*) with respect to the curvature of the front (≈ 750*μm*). Notably, λ = 45 μm, parameterized by matching experimentally observed sector shapes, is remarkably consistent with the *T* = 25 μm found for the Interface-growth model under the predicted relation *T* = λ/2 (see SI Fig. 11 and 10).

Our results so far suggest that slower growing mutant clones at the advancing frontier are mechanically screened from the competition with wild-type cells, an effect that is strongest for small clones. Clones emanating from single pre-existing or newly arisen mutant cells should therefore benefit the most and exhibit a nearly neutral lineage dynamics. To test this prediction, we inoculated a small fraction of single mutant cells interspersed into a background of wild-type cells. These experiments show that mutant clones form elongated streaks similar to what we see in the Interface-growth model (compare Fig. 4a and b). Moreover, the probability of a clone to remain at the front decays several orders of magnitude more slowly than the null expectation, which ignores surface tension (Fig. 4c).

The inefficient purging of slow growing mutants can have severe evolutionary consequences when the selection pressure suddenly shifts in favor of the mutants. For instance, costly drug resistant mutants may persist long enough to trigger resurgent growth upon application of the corresponding drug [13, 1]. To test whether the screening of fitness effects promotes such an evolutionary rescue [2, 5], we first grew a colony harboring a small fraction of single mutant cells, as described above. These mutants, while having a growth rate deficit with respect to the wild-type cells in the absence of the drug, are resistant to the antimicrobial Hygromycin B. Despite their initial growth disadvantage, a substantial fraction of mutant clones persisted at the front even after 4 days of growth (Fig. 4e). We then subjected the entire colony to high levels of Hygromycin B, completely stalling wild-type growth. While mutant clones trapped inside the colony remained effectively confined by surrounding wild-type cells [12], the clones persisting at the leading edge radiated out establishing an all-mutant population expansion (Fig. 4f).

The crowding-induced screening of fitness effects should also affect small clones of *beneficial* mutations in the early stages when they are still small. We therefore grew yeast colonies of wild-type cells mixed with a low fraction of faster growing mutants (Fig. 4g). The mutants that were to establish generated clones exhibiting a characteristic funnel-like shape, with a filament-like beginning and wide opening towards the end, similar to previous observations [19, 32]. The slow initial part indicates a prolonged time for mutant establishment, which is typically associated with a reduced establishment probability [14, 9].

In summary, our results demonstrate that evolutionary processes have a mechanical basis in crowded populations. Growth-induced pushing forces generate group-level interactions that screen selective differences due to an effective surface tension. Compared to non-mechanical models, selection is less efficient in weeding out deleterious mutations, which can be rescued by an environmental change. Mechanical screening effects merely require steric interactions and could therefore substantially affect the evolutionary dynamics in any crowded cellular population, such as biofilms, multi-cellular tissues or tumors [38, 22, 21], by increasing clonal interference [17, 25], promoting the accumulation of dele-terious mutations [23, 27, 16, 4], or facilitating the rescue of costly phenotypes by compensatory mutations [40, 1].

**Figure 4:**
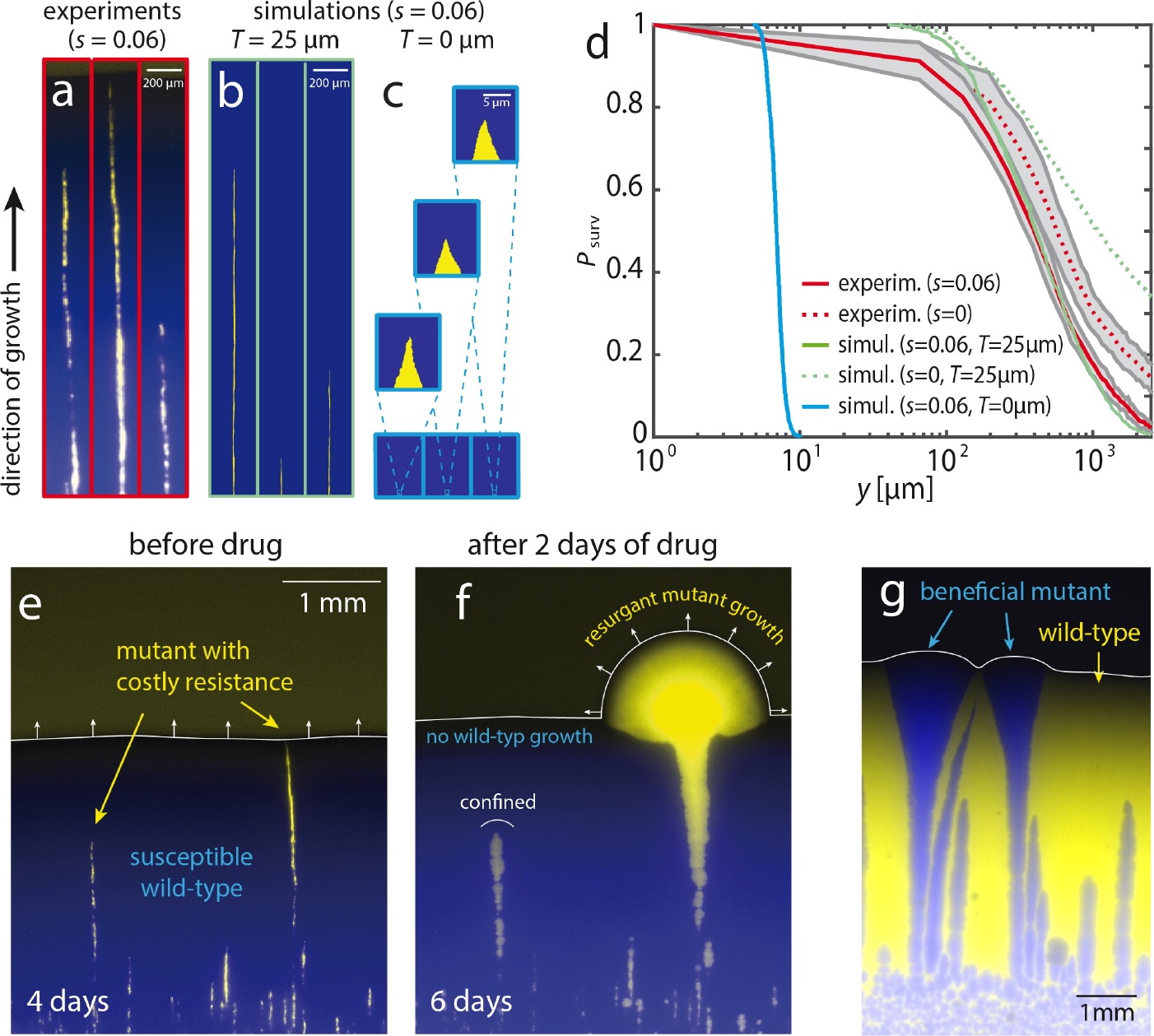
Slow purging of costly drug resistance mutations can lead to resurgent growth upon drug application. a) Fluorescent micrographs of thin sectors originating from single cells of a slower growing mutant phenotype (yellow, *s* = 0.06) interspersed in a wild type colony (blue). Images show three characteristic mutant lineages that are extended in the direction of motion. b) To compare this scenario to our simulations, we added boundary drift to the Interface-growth model used in Fig. 2. This stochastic Interface-growth model (boundary diffusion coefficient *D* = 0.025 μm) exhibits similarly long streaks. c) Simulations akin to those in b) but void of surface tension. In this scenario, equivalent to the Local-speed model with boundary drift, mutants are purged extremely quickly. The zoomed-in insets are magnified by a factor of 32. d) Fraction of clones still present at the front, *P*_surv_, as a function of front progression y for experiments, stochastic Interface-growth model simulations (*T* = 25 μm) and the stochastic Local-speed model (*T* = 0) (SI Sec. 5.2). Experimental curves are averages of 12 independent colony fronts and the shaded gray area indicates the standard deviation. For neutral experiments (*s* = 0) mutant cells are substituted for cells with a growth rate identical to that of the wild-type. Both, the experimental observations and the Interface-growth model are much closer to their respective neutral scenarios than to the rapidly decaying null expectation. e) The mutant strain (yellow) is resistant to the antimicrobial Hygromycin B. f) Application of this drug to the entire colony completely arrests wild-type growth. As a result, any resistant mutant still present at the front (even if just as one single cell) will rapidly expand and lead to all-mutant resurgent growth. Resistant clones within the colony remain confined g) Beneficial mutant clones have a funnel-like transience prior to establishment, an inverse of the deleterious funnel in Fig. 1c. The slight broadening of faster growing clones deep within the colony is due continued to out-of-plane growth.

